# CSOA: A Novel Single-Cell Gene Set Enrichment Analysis Method with Comprehensive Benchmarking

**DOI:** 10.64898/2026.07.16.738892

**Authors:** Andrei-Florian Stoica, Kejun Yao, Jie Wang, Xueli Xu

## Abstract

Single-cell gene set enrichment analysis is widely used to evaluate the activity of gene sets in individual cells, as measured by single-cell sequencing technologies. However, existing methods often generate ambiguous scores that cannot reliably distinguish cells enriched for a biological signal from background cells. To address this limitation, we developed Cell Set Overlap Analysis (CSOA), a novel method for gene set enrichment analysis that leverages gene pair relationships by quantifying pairwise overlaps between high-expression cell sets constructed for each signature gene. We benchmarked CSOA against sixteen established methods representing five methodological classes: direct scoring, rank-based scoring, model-based scoring, matrix decomposition, and overrepresentation analysis. Our evaluation framework introduces novel metrics tailored for the gene set scoring problem, such as score coverage and silhouette rank alignment. They are used alongside traditional metrics for binary classification, such as the Matthews correlation coefficient and area under the receiver operating characteristic (AUROC). CSOA showed superior accurate annotation of cell types and specific biological processes compared with competing approaches. This advantage was particularly pronounced in the class boundary determination benchmark, where it ranked the first in all evaluated datasets. CSOA also outperformed most of the compared methods in computational efficiency. Notably, CSOA’s combination of outstanding performance in the score coverage metric and solid overall performance positions it as a uniquely well-suited method for distinguishing cells enriched for specific biological signals.

## Introduction

Single-cell gene set enrichment analysis (scGSEA) has been widely employed to identify enriched biological functions in single-cell RNA sequencing (scRNA-seq) data [1]. Typically, the gene sets represent biological signals, such as cell types, biological processes, or signaling pathways [2], enabling scGSEA methods to assess the enrichment of functional modules at single-cell resolution. Initially, scGSEA methods were adapted from tools developed for bulk RNA-seq data. More recently, scGSEA methods have been specifically developed for scRNA-seq data to better handle high dropout rates and technical noise, while also providing greater scalability for large numbers of cells [3].

Despite these advancements, several challenges remain, particularly in effectively prioritizing genes within the input gene set. Since not all genes contribute equally to characterizing a biological signal, an ideal scGSEA method should automatically infer the relative importance of each gene without relying on explicit prior weights. Another key challenge is achieving a clear separation between cells enriched for the biological signal and background cells. Existing scGSEA methods often fail to clearly separate cells with high biological signal scores from low-scoring cells. Instead, they produce intermediate scores suggestive of mixed or transitional populations, making it difficult to robustly identify cells characterized by strong biological signals.

Here, we present Cell Set Overlap Analysis (CSOA), a novel scGSEA method designed to address these challenges. Unlike traditional methods that aggregate gene expression values, CSOA leverages pairwise overlaps among high-expression cell sets constructed for each signature gene. Specifically, for each gene, CSOA identifies the cells with high expression and then computes pairwise overlaps among these cell sets to compute enrichment scores. This overlap-based approach naturally captures gene co-expression patterns while automatically weighing genes according to their discriminative power. A key feature of CSOA is that pairwise overlaps are sufficient to achieve great performance while avoiding the computational burden that higher-order overlaps would impose.

We benchmarked CSOA against 16 established methods spanning five methodological categories: direct scoring (AddModuleScore [4], SiPSiC [5], and Zscore [6]), rank-based scoring (AUCell [7], GSVA [8], JASMINE [9], Singscore [10], ssGSEA [11], and UCell [12]), model-based scoring (MDT, MLM, ULM [13], and VAM [14]), matrix decomposition (Pagoda2 [15] and PLAGE [16]), and overrepresentation analysis (ORA [13]). We systematically evaluated the performance of all methods across three complementary dimensions: class boundary determination, Matthews correlation coefficient (MCC), and global ranking consistency.

Computational time and memory usage were also assessed for each method. CSOA demonstrates superior accuracy compared with competing methods while providing substantial improvements in computational speed and memory efficiency.

## Methods

### The CSOA algorithm

First, CSOA constructs a cell set for each signature gene by identifying the set of cells with gene expression above the 90th percentile, calculated across all cells expressing that gene. P-values for all pairwise overlaps between these cell sets are computed using the hypergeometric test via the phyper function from the R stats package [17]. For two cell sets of sizes *n_i_* and *n_j_* selected from a dataset with *N* cells, with an intersection of size *k_ij_*, the p-value is given by:

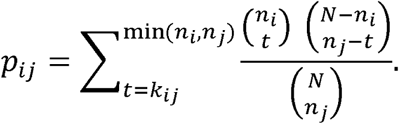

These p-values are adjusted using the Benjamini-Yekutieli correction method for multiple testing via the p-adjust function from the stats package (method = “BY”) [18]. Only overlaps with an adjusted p-value < 0.05 are retained for subsequent analysis.

Second, the retained overlaps are ranked. Each overlap receives two preliminary ranks: one based on its adjusted p-value 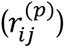 and another based on its shared cell ratio 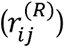. The shared cell ratio is defined as the observed overlap divided by the expected overlap under the hypergeometric null model:

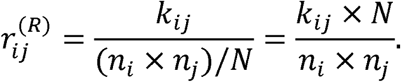

Each overlap is interpreted as an edge (*i*, *j*) ∈ *E* in a graph *G* = (*V*, *E*), where genes correspond to vertices *V*. The two initial ranks are then refined based on graph connectivity. Specifically, for each overlap, both the p-value rank and the cell ratio rank are updated to the minimum of their current rank and the lowest rank of any adjacent edge. Denoting the updated ranks as 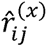 where x ∈ {p,R} indicates the rank type:

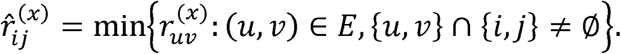

The final rank (called the aggregate rank) for each overlap is determined by averaging these two updated ranks, with ties resolved by selecting the minimum rank. Top overlaps are selected using a rank cutoff defined as the most frequent rank when unique, or as the average of the minimum and maximum ranks among the most frequent ranks. Overlaps with ranks at or below this cutoff are retained for scoring, while others are discarded.

Finally, the selected overlaps are assigned weights based on their aggregate ranks. Let {*q*_1_, *q*_2_, …, *q_L_*} denote the *L* unique ranks among the top overlaps, sorted in ascending order. To generate weights from ranks, a sequence of *L* + 1 values linearly decreasing from e to 1 is generated, and weights are assigned by applying the natural logarithm. Thus, the weight (*w*_ℒ_) corresponding to rank index ℒ is:

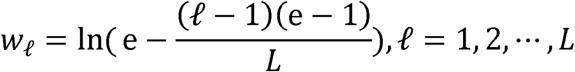

where e denotes Euler’s number (e ≈ 2.718). The edge weight *w_ij_* for each selected overlap (*i*, *j*) is the weight *w*_l_ associated with its aggregate rank.

Cell-level scores are then computed. Each overlap contributes a gene pair score 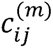 to each cell *m*, computed as the product of its edge weight (*w_ij_*) and the min-max-normalized expression levels 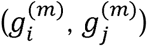 of the two genes:

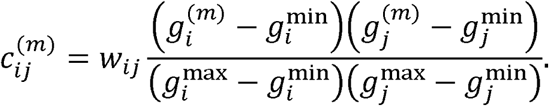

The aggregate score (*C_m_*) for each cell is the sum of contributions from all selected overlaps:

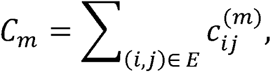

and final cell scores (*Ĉ_m_*) are obtained through min-max normalization:

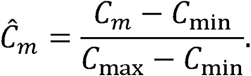

### scRNA-seq dataset preprocessing

Four human scRNA-seq datasets were analyzed using R (version 4.5.2). The Baron pancreas dataset [19] was loaded via the scRNAseq package [20], while the remaining three datasets were loaded from the PangliaoDB database [21]. Dataset details are summarized in Table S1.

All datasets were processed using the Seurat package [22]. The Baron dataset, originally provided as a SingleCellExperiment object, was converted to a Seurat object using the as.Seurat function. For each dataset, genes expressed in fewer than 10 cells were filtered out prior to standard Seurat preprocessing, which included normalization, variable feature selection, data scaling, and principal component analysis (PCA). Additional quality control was applied to two datasets: for the Merkel cell carcinoma dataset, the cells with a percentage of mitochondrial genes ≥ 10% were removed; for the peripheral blood mononuclear cells (PBMC) dataset, cells with a percentage of mitochondrial genes ≥ 5% were removed. Mitochondrial gene percentage was regressed out during scaling using the ScaleData function.

Dimensionality reduction and visualization were performed using Uniform Manifold Approximation and Projection (UMAP). The number of PCA dimensions used as input for UMAP was determined as the lowest dimension at which the difference in explained variance between consecutive principal components dropped below 0.1%. Clustering was performed on the Merkel cell carcinoma and PBMC datasets. First, the FindNeighbors function from Seurat was run using the two UMAP dimensions as input. Then, clustering was performed with the FindClusters function from Seurat at the resolution of 0.4 for the Merkel cell carcinoma dataset and 0.1 for the PBMC dataset. The clusters were merged based on functional identities, which were determined by assessing the top entries returned by the enrichGO function from the clusterProfiler package [23] with cluster markers identified by the FindMarkers function from Seurat as input.

### Gene set construction

In this study, four scRNA-seq datasets were analyzed. The pancreas and lung datasets were used for cell type identification, whereas the Merkel cell carcinoma and PBMC datasets were used to identify functional identities. For the pancreas and lung datasets, marker gene sets for each cell type were obtained from PanglaoDB. Genes absent from the datasets were excluded.

For the Merkel cell carcinoma and PBMC datasets, gene sets were constructed through gene ontology (GO) enrichment analysis using enrichGO with cluster markers as input. Each gene set comprised cluster markers from the enriched GO term defining the cluster identity.

### Evaluation metrics for benchmarking scGSEA methods

The benchmark included three types of evaluation metrics: class boundary determination, global evaluation, and MCC. We included MCC as a distinct evaluation metric because it provides a robust and balanced measure for binary classification tasks [24].

Class boundary determination metrics evaluate whether cells exhibiting strong enrichment signals for the input gene set are effectively separated from other cells. These metrics are computed at each possible separation point (the unique scores for each scGSEA method), after which the average metric score is calculated. The separation point with the highest average metric score is taken as the cutoff delineating high method scores, indicating significant enrichment. The highest average metric score (converted to percentage) is further referred to here as the boundary benchmark score.

These metrics include four traditional metrics (sensitivity, specificity, precision, and accuracy), as well as two novel ones: size proximity and score coverage. All are defined based on true positives (*TP*), true negatives (*TN*), false positives (*FP*) and false negatives (*FN*):

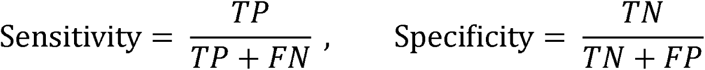

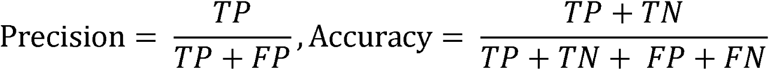

Size proximity is used to evaluate how balanced the predicted class sizes are relative to the ground truth and is defined as follows:

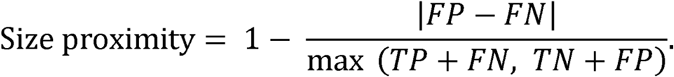

For score coverage, we employ the per-cell score *δ_m_* calculated by scGSEA methods (e.g., the score *Ĉ_m_* from CSOA). We then compute:

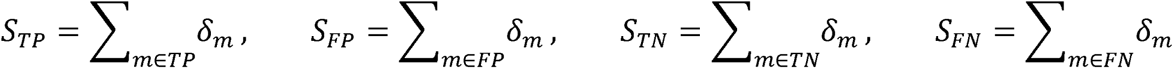

Score coverage is then defined as:

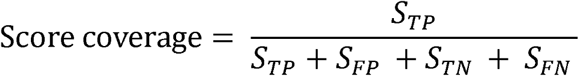

This metric quantifies the proportion of total scores attributable to true positive cells, reflecting whether the model allocates higher scores to biologically meaningful cells.

The global evaluation metrics assess whether the overall distribution of predicted scores aligns with the ground truth. They provide threshold-independent assessments of prediction quality, directly returning a single value without requiring the maximization of an average of multiple metrics. These metrics include the area under the receiver operating characteristic (AUROC), the area under the precision-recall curve (PRAUC), label Jaccard score and label cosine score, as well as three novel metrics: label rank alignment, silhouette rank alignment and centrality. We will next introduce the novel metrics.

Let ***f***, ***y*** ∈ ℝ*^n^* be two real-valued vectors (e.g., predicted scores and ground-truth labels). Let *ρ_f_*(*i*) and *ρ_y_*(*i*) denote the ranks of element *i* in ***f*** and ***y***, respectively (*i* = 1, …, *n*). The min-max-normalized ranks are:

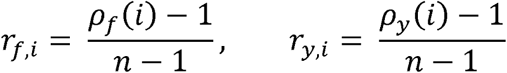

Let *r_f_*_,(1)_ ≤ *r_f_*_,(2)_ ≤ ⋯ ≤ *r_f_*_,(*n*)_ denote the sorted values of {*r_f_*_,*i*_}, and similarly for {*r_y_*_,*i*_}. Then rank alignment is defined as:

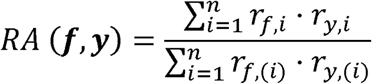

Label rank alignment is then defined as the rank alignment of the vector of predicted scores and the binary vector determined by ground-truth identities.

Next, to define the notion of silhouette rank alignment, we will introduce the concept of normalized silhouette. We will first provide the definition of silhouette [25]. Let *Z* = {*z*_1_, *z*_2_, …, *z_n_*} be a dataset (e.g., UMAP coordinates) partitioned into *K_clust_* clusters 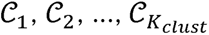. For each data point *z_i_* ∈ *C_j_*, define the average distance between the point and all other points in the same cluster as follows:

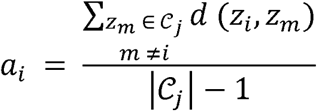

Define the minimum average distance between the point and the points in other clusters as follows:

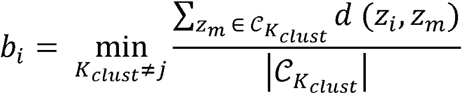

The silhouette coefficient *s_i_* for point *z_i_* is defined as:

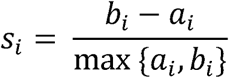

We define the normalized silhouette as a matrix with *K_clust_* columns, each corresponding to a cluster. We will now detail the procedure of calculating normalized silhouette for cluster *C_j_*. First, all cells outside of the cluster will be assigned a normalized silhouette value of 0. Then, let *A* = {*s_i_* ∣ *z_i_* ∈ *C_j_*} be the set of silhouette values for in-cluster cells. We first define a fictitious lowest silhouette value to prevent assigning a normalized silhouette score of 0 for any in-cluster cell: *s*_fict_ = 2 × min(*A*) − *s*_(2)_, where *s*_(2)_ is the second-lowest value of *A*. Next, we perform min-max normalization over the extended set *A* ∪ {*s*_fict_}:

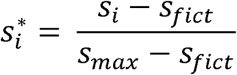

We can now define silhouette rank alignment as the rank alignment of the vector of predicted scores and the column corresponding to the cluster in the normalized silhouette matrix.

The next metric, centrality (*μ*), also employs the normalized silhouette. We first define the center of mass of a scGSEA method score vector 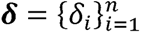, where *n* represents the number of cells, and ***z****_i_* represents the UMAP coordinates of cell *i*:

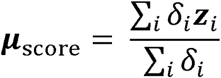

The center of mass of normalized silhouette for cluster *C_j_* is defined as follows:

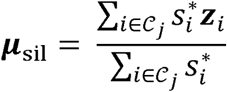

We are now ready to define centrality:

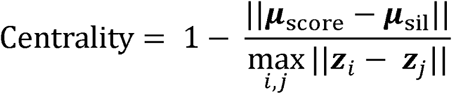

Finally, we introduce two global metrics that depend upon the results obtained in the class boundary benchmark assessment: label Jaccard score and label cosine score. These metrics first convert the scGSEA method scores into binary predictions based on the boundary obtained for each method in the class boundary benchmark assessment. Then, the Jaccard similarity and the cosine similarity between the binary predictions and the truth labels are computed. Thus, let *P_i_* be the predicted label for cell *i*, and *T_i_* denote the true label. The two metrics are computed as follows:

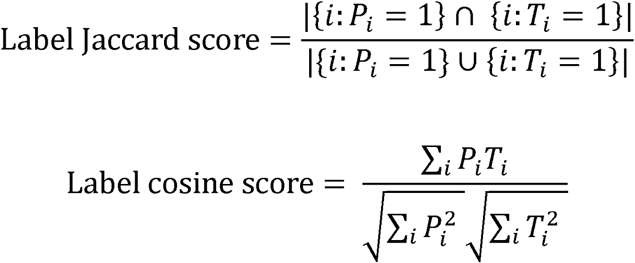

The MCC defined below has two implementations. The first, comprehensive MCC, is used as a class boundary determination metric, where values are calculated at each separation point and the maximum value is retained. The second, direct MCC, is a global metric computed based on the results from the class boundary determination metric.

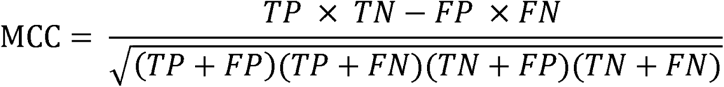

## Results

### Overview of CSOA

Here, we provide an overview of the CSOA workflow, which consists of six major steps (Figure 1 and Figure S1). Starting from an scRNA-seq dataset and a gene set, CSOA generates per-cell gene set scores. First, CSOA constructs high-expression cell sets for each gene in the signature. Second, the significance of cell set overlaps is assessed using hypergeometric tests. Third, two ranks are built based on the adjusted p-value of the overlap and the shared cell ratio of the observed and expected number of cells assuming a hypergeometric model, and the ranks are updated based on graph connectivity and averaged to obtain the final rank. The cutoff for retaining the top overlaps is determined based on the most frequent rank. Fourth, overlap weights ranging from 0 to 1 are calculated by taking the natural logarithm of values between 1 and e corresponding to distinct ranks. Fifth, gene pair scores are then computed for each cell by multiplying overlap weights with the min-max-normalized expression of each involved gene. Finally, gene pair scores are summed for each cell and min-max-normalized to generate the final CSOA scores.

**Figure 1:**
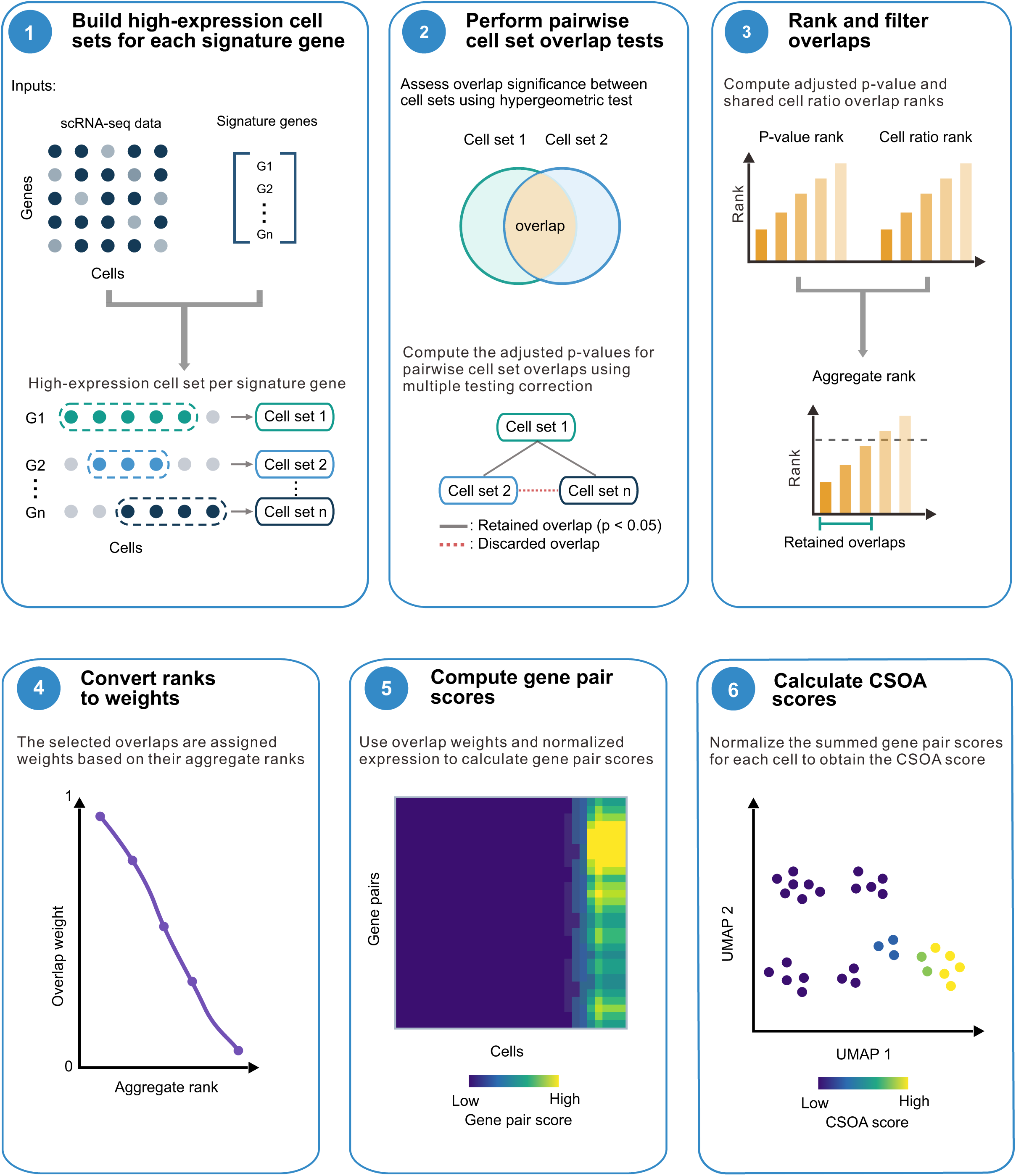
Overview of CSOA. CSOA includes six main steps: (1) Construction of high-expression cell sets for each signature gene. (2) Statistical assessment of pairwise cell set overlaps using hypergeometric tests. (3) Ranking the overlaps based on adjusted p-value and shared cell ratio. (4) Computation of overlap weights by assigning logarithmically decreasing values from 1 to 0 to aggregate ranks. (5) Computing gene pair scores for each cell by multiplying overlap weights with the min-max-normalized expressions of the two involved genes. (6) Construction of CSOA scores by summing all gene pair scores for each cell and subjecting these scores to min-max normalization.

### CSOA accurately and reliably annotates cell types

On the Baron pancreas dataset, CSOA scores for the acinar gene set strongly correspond to acinar cells (Figure 2A), with a boundary benchmark score (defined in Methods) of 95.41% (Figure 2B). For the ependymal cells in the lung proximal airway stromal cells dataset, the boundary benchmark score was 98.99% (Figure 2C–D). For the more challenging task of identifying clusters based on biological functions from gene ontology, CSOA also achieved an accurate annotation of biologically defined clusters. For instance, CSOA obtained a boundary benchmark score of 94.98% for the chromosome segregation gene set in the Merkel cell carcinoma dataset (Figure 2E–F). Similarly, CSOA obtained a high boundary benchmark score (92%) for the cell killing gene set in the PBMC dataset (Figure 2G–H).

**Figure 2:**
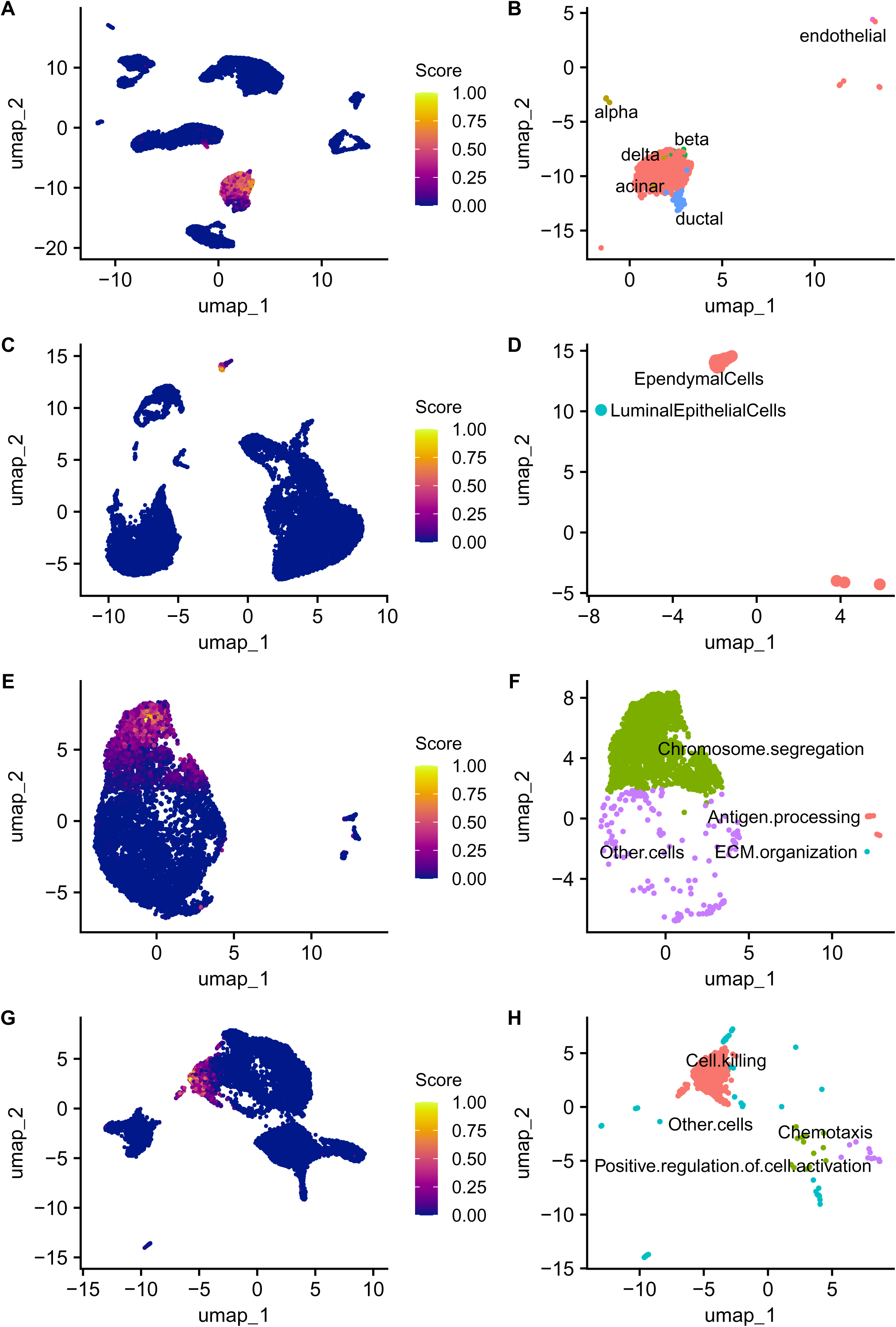
Examples of CSOA prediction. CSOA scores and binary predictions obtained using: the acinar gene set on the Baron pancreas dataset (A, B); the ependymal gene set on the lung proximal airway stromal cells dataset (C, D); the chromosomal segregation gene set on the Merkel cell carcinoma dataset (E, F); the cell killing gene set on the peripheral blood mononuclear cells (PBMC) dataset (G, H).

To assess the reliability of CSOA for single-cell gene set enrichment analysis, we evaluated how prediction performance depends on gene signature quality. Specifically, a fixed fraction of genes was removed from each gene signature (gene loss) or replaced with randomly selected genes (noise) to examine prediction accuracy. All combinations of 20%, 50% and 80% gene loss and noise were assessed, and CSOA delivered generally accurate results when gene loss and noise were set to 50% or below (Figure S2–S4). These findings indicate that CSOA can reliably identify biological signals.

### CSOA scoring is highly divergent from existing methods

Given that CSOA employs a novel gene pair-based strategy for scGSEA, we compared it with 16 existing scGSEA methods to determine whether CSOA exhibits distinct performance. When per-cell scores are replaced with percentages of the total score, CSOA occupied a distinctly isolated position in the embedding space across all four datasets, clearly separated from the dense cluster formed by most of the methods (Figure S5). In contrast, when methods were evaluated based on binary predictions, CSOA showed high similarity to other high-performing methods such as PLAGE (Figure S6), registering Jaccard similarity scores of 0.86, 0.9, 0.86 and 0.78 with the latter method.

### CSOA outperformed the other methods in class boundary determination

Next, we benchmarked CSOA against 16 scGSEA methods using a comprehensive set of class boundary determination metrics, including sensitivity, specificity, precision, accuracy, size proximity, and score coverage. Size proximity quantifies how closely the number of cells predicted to belong to a given identity matches the ground truth, whereas score coverage measures the proportion of the total method-derived score attributed to cells of the identity of interest. Across all four datasets, CSOA consistently achieved the highest performance among the 17 evaluated scGSEA approaches according to the class boundary determination benchmark (Figure 3). Notably, CSOA demonstrated a particularly strong advantage in score coverage, together with UDT and MDT (Figure S7).

**Figure 3:**
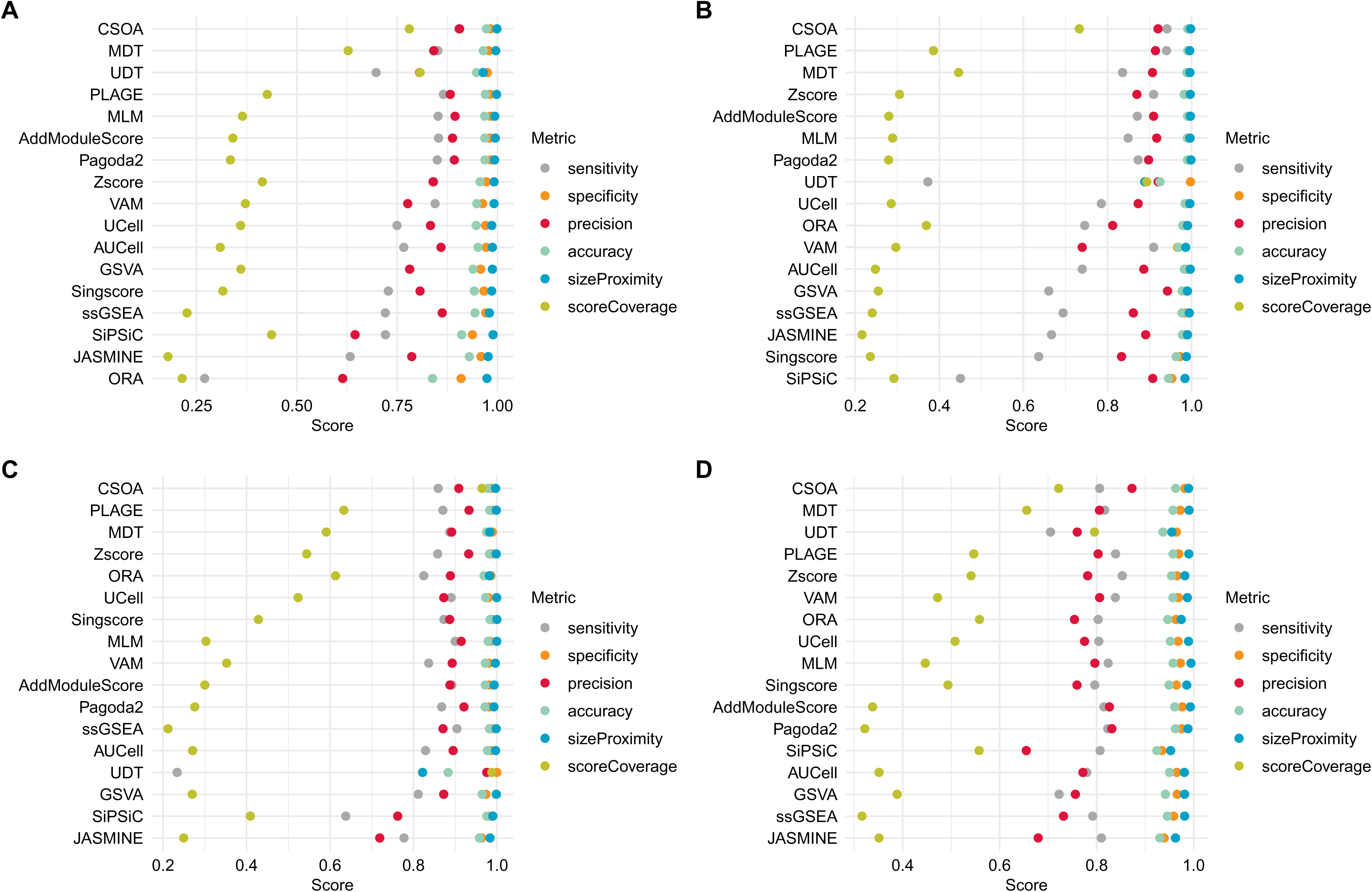
Class boundary determination benchmark. CSOA achieved the best result in the class boundary determination benchmark on the Baron pancreas (A), the lung proximal airway stromal cells (B), the Merkel cell carcinoma (C), and the PBMC (D) datasets, with the score coverage metric showcasing excellent performance.

### CSOA shows superior performance on the Matthews correlation coefficient metric

We then assessed the performance of CSOA and other methods on the MCC metric which evaluates the overall performance of binary classification. In the comprehensive MCC assessment, CSOA ranked first on the Baron pancreas dataset, showing significant improvements over all other methods, particularly for the identification of delta cells (Figure 4A). CSOA likewise achieved the top rank on both the lung proximal airway dataset (Figure 4B) and the PBMC dataset (Figure 4D). In contrast, CSOA ranked 9th on the Merkel cell carcinoma dataset (Figure 4C). Highly similar results were observed in the direct MCC assessment (Figure S7).

**Figure 4:**
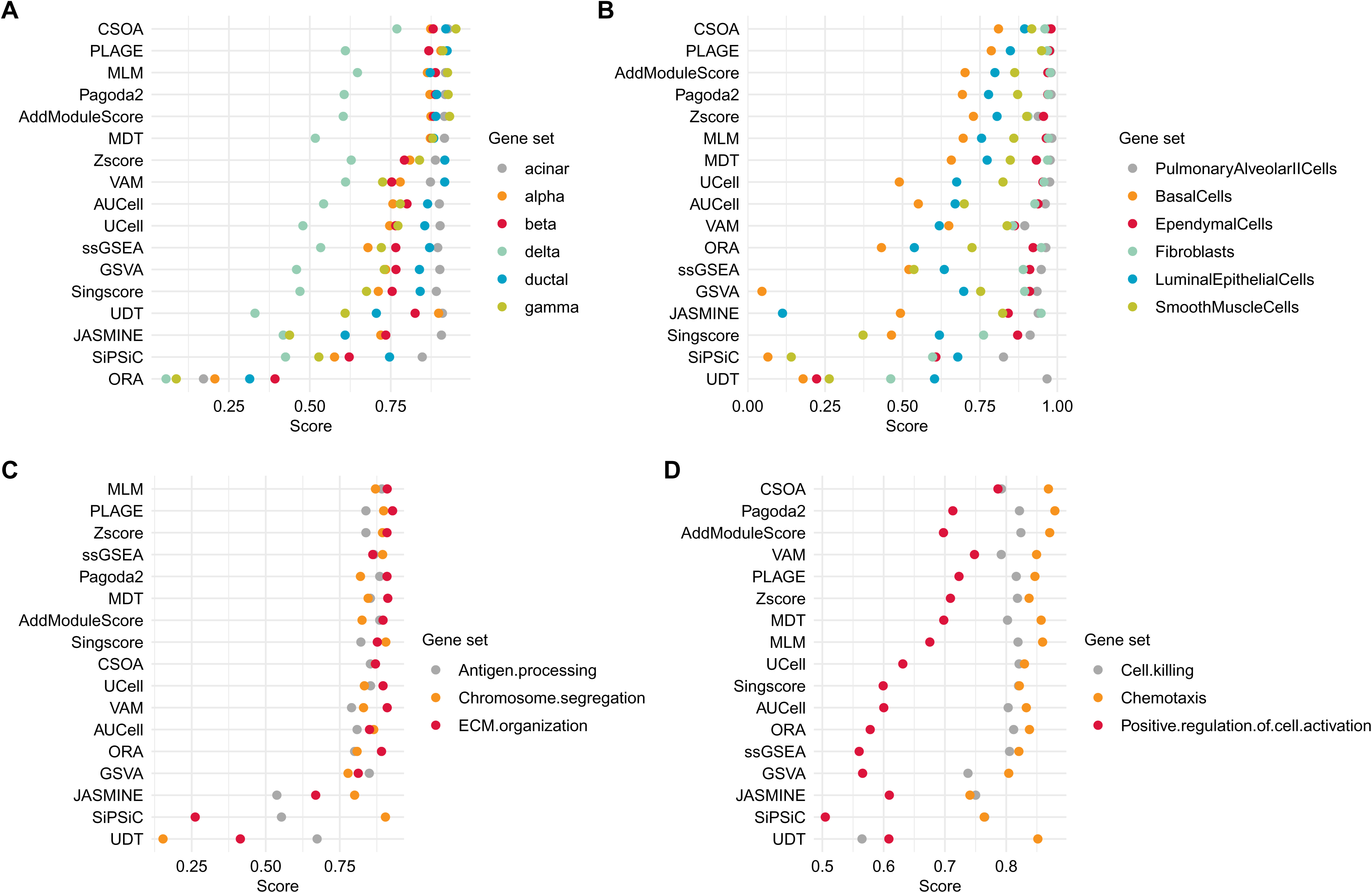
Comprehensive MCC benchmark. In the comprehensive MCC assessment, CSOA ranked the first on the Baron pancreas (A), lung proximal airway stromal cells (B), and PBMC (D) datasets, and the ninth on the Merkel cell carcinoma dataset (C).

### CSOA shows good performance in the global evaluation benchmark

We further employed the global evaluation to determine whether the overall distribution of predicted scores aligns with the ground truth. The global evaluation benchmark shows similar patterns with those observed in the MCC benchmark, with CSOA displaying strong performance for cell type-associated gene sets but comparatively lower performance for gene sets representing biological functions. CSOA achieved the best overall performance among all evaluated scGSEA methods on the Baron pancreas dataset (Figure 5A) and the lung proximal airway stromal cells dataset (Figure 5B), but ranked the 9th on the Merkel cell carcinoma dataset (Figure 5C) and the 8th on the PBMC dataset (Figure 5D). Among the global evaluation metrics, CSOA showed superior performance in centrality, which assesses the proximity between the center of mass of CSOA scores and that of the normalized silhouette of the cluster of interest (Figure S3).

**Figure 5:**
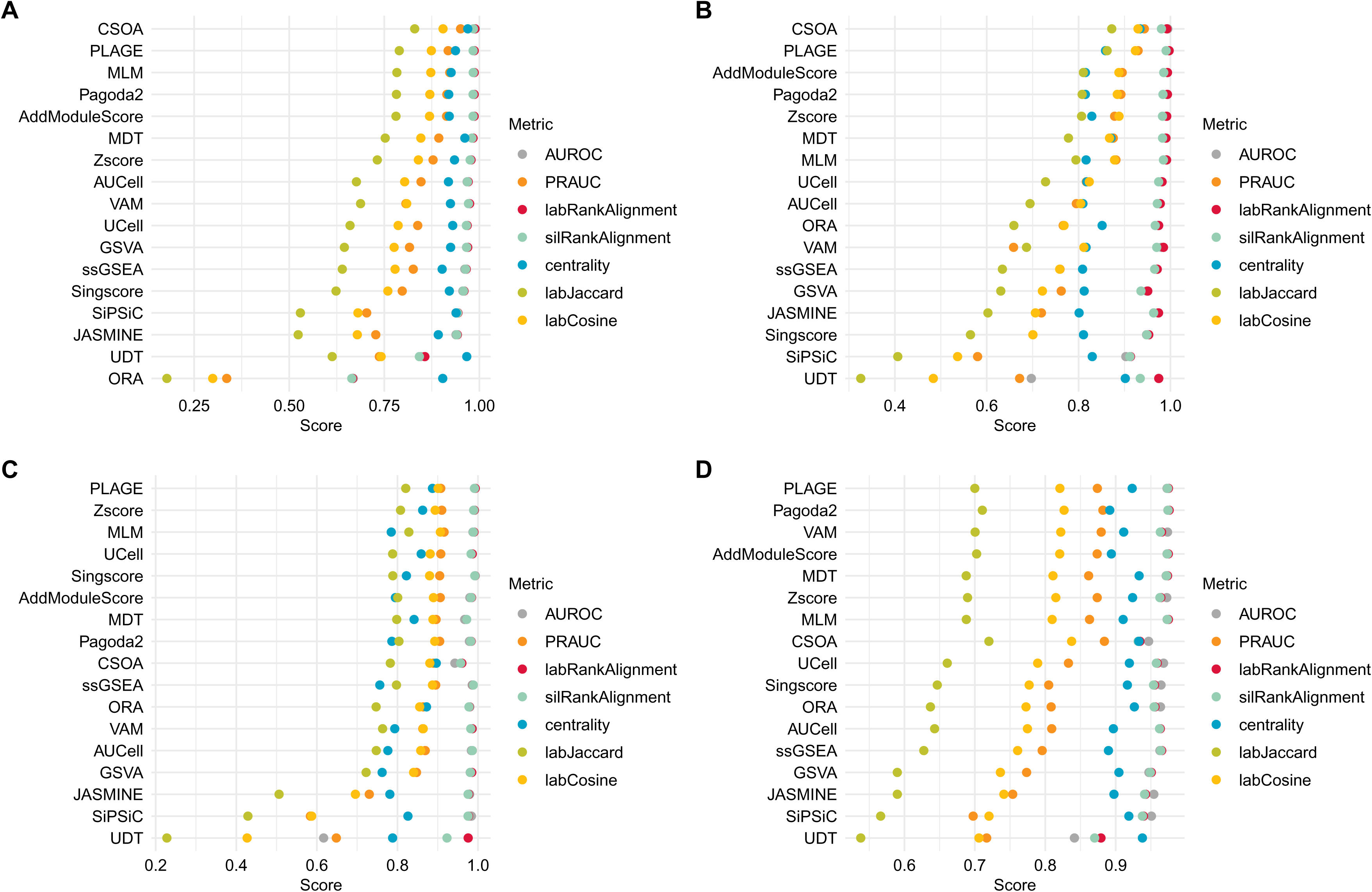
Global evaluation benchmark. In the global evaluation benchmark, CSOA ranked the first on the Baron pancreas (A) and lung proximal airway stromal cells (B) datasets, the ninth on the Merkel cell carcinoma dataset (C), and the eighth on the PBMC dataset (D).

### CSOA is efficient in terms of time and memory usage

To systematically compare the computational efficiency of CSOA with other methods, we calculated the time and memory usage separately across four datasets. CSOA was the third-fastest method on the Baron pancreas (Figure 6A) and Merkel cell carcinoma (Figure 6C) datasets, and the fourth-fastest method on the lung proximal airway stromal cells (Figure 6B) and the PBMC (Figure 6D) datasets. In terms of memory efficiency, CSOA ranked fifth in each of the four datasets (Figure 7).

**Figure 6:**
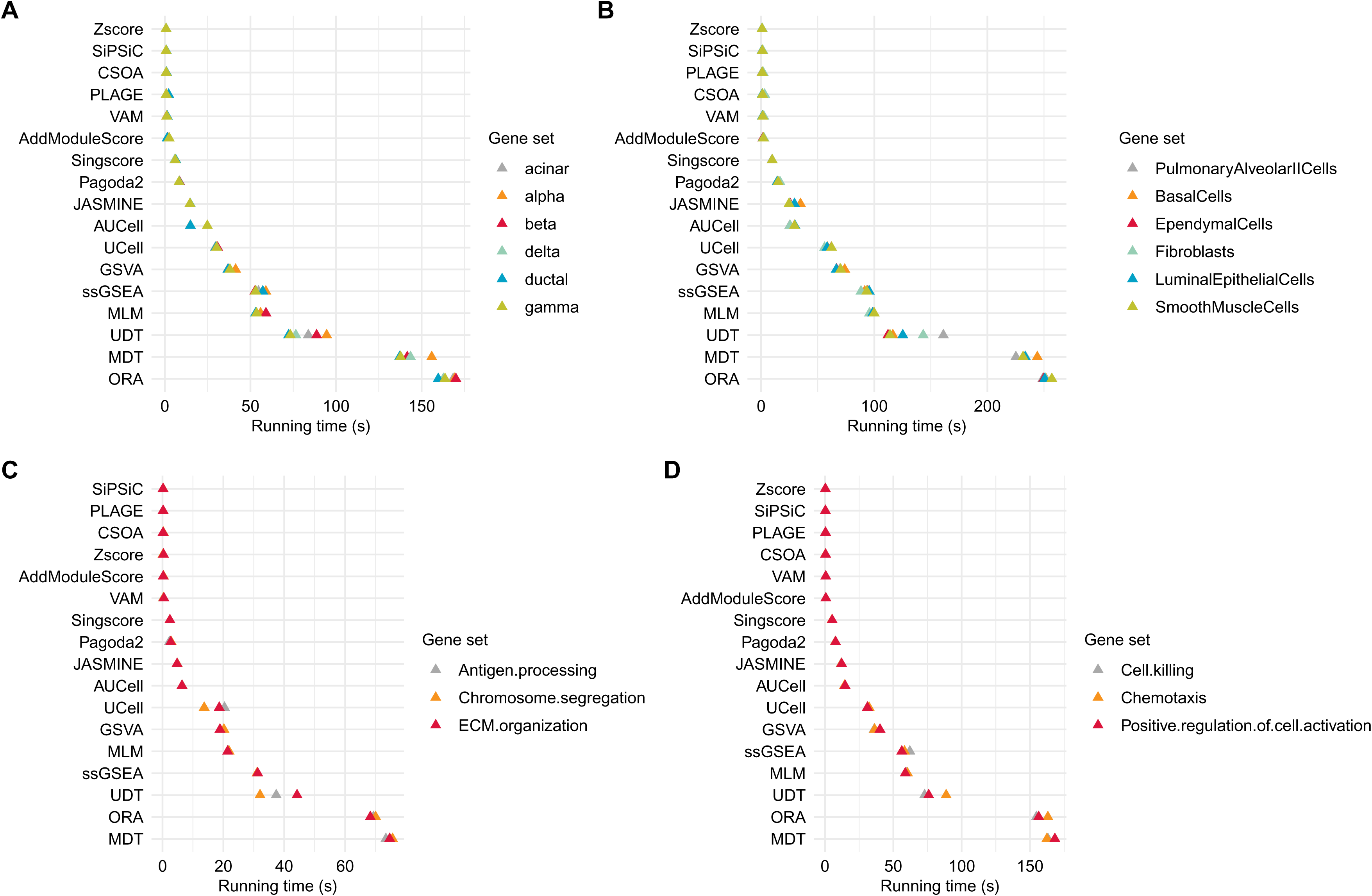
Elapsed time benchmark. CSOA was the third-fastest method on the Baron pancreas (A) and Merkel cell carcinoma (C) datasets, and the fourth-fastest method on the lung proximal airway stromal cells (B) and PBMC (D) datasets.

**Figure 7:**
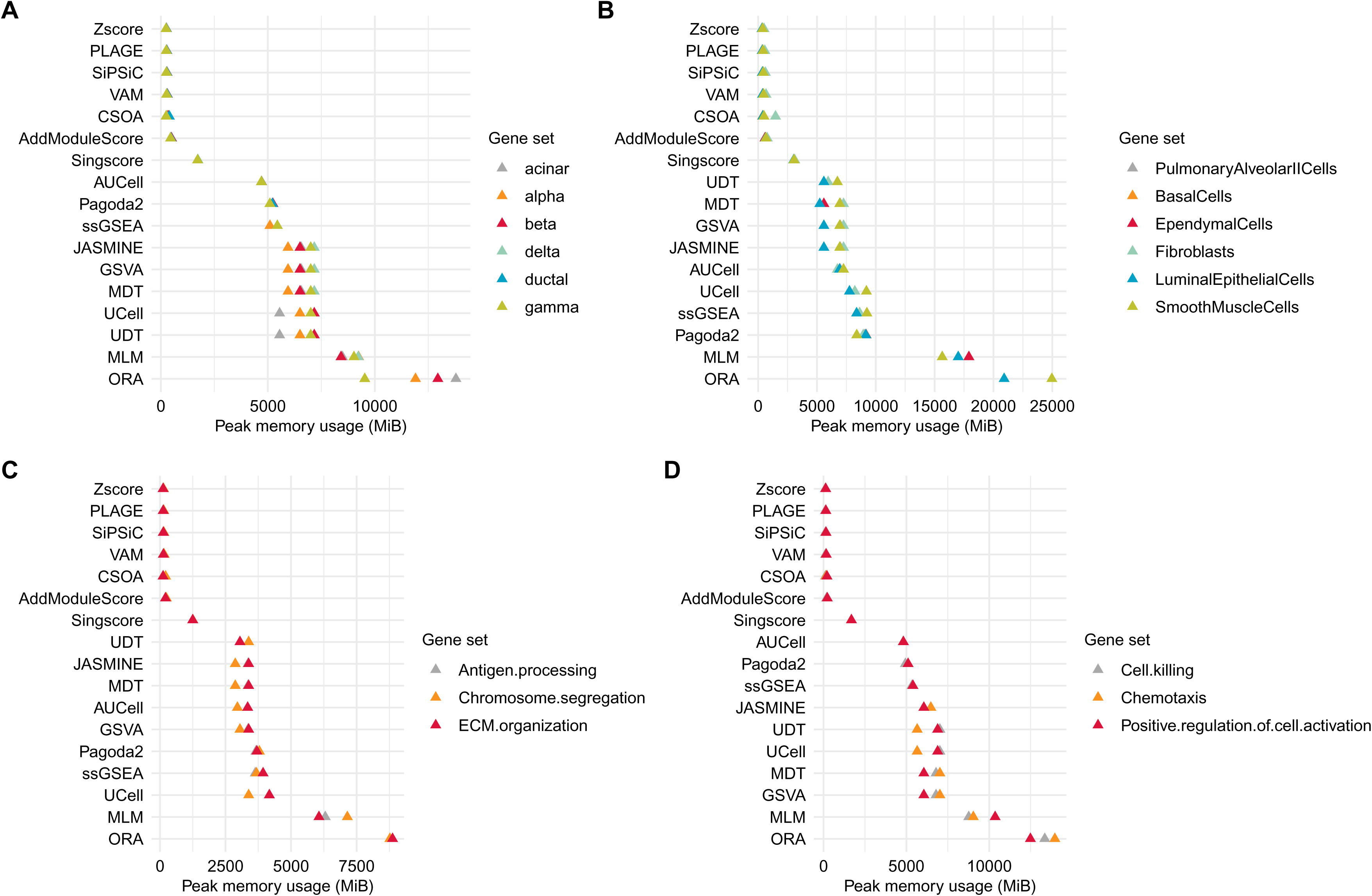
Memory usage benchmark. CSOA registered the fifth-lowest peak memory usage on the Baron pancreas (A), lung proximal airway stromal cells (B), the Merkel cell carcinoma (C), and the PBMC (D) datasets.

## Discussion

Currently, scRNA-seq technologies have been widely applied to characterize cellular heterogeneity at unprecedented precision and depth. However, the frequent problem of scoring a gene set of biological interest still requires new methodological approaches. Although numerous scGSEA methods have been developed, none of them became established as a gold standard. Comprehensive benchmarking studies evaluating a sufficiently large number of scGSEA methods remain lacking, with recent efforts typically focusing on only a limited number of methods [26, 27]. Consequently, researchers often lack sufficient information to select an appropriate scGSEA method for a given application. Because scGSEA methods vary considerably in their scoring approaches and enrichment calculations, this gap limits the generalizability of reported findings.

This paper introduces CSOA, a novel scGSEA method that innovates upon existing methods through focusing on gene pairs rather than individual genes, and compares CSOA to 16 other scGSEA methods by using both traditional and novel metrics. CSOA achieves substantially better separation of cell populations characterized by the biological signal of interest from those lacking it, as evidenced by consistently robust performance in the score coverage metric across diverse datasets and gene sets.

The development of CSOA also raises questions of theoretical interest. One question concerns the similarity between the results obtained by CSOA and those obtained by decision tree-based methods (MDT and UDT) on the score coverage and centrality metrics. Exploring the mechanisms driving this similarity may open the way for future improvements in the problem of gene set scoring. Another potential avenue for future research is a consensus model integrating the scores generated by the top-performing scGSEA methods.

However, CSOA does have some limitations. Generally, CSOA performed better for cell types than for functional identities, which may be explained by the fact that in practice, functional identities, unlike cell types, are often not truly specific to a single cluster but are partly shared with neighboring clusters. Furthermore, despite its strong performance in class boundary determination, CSOA registered lower scores in the global evaluation benchmark, partly due to its performance on the label alignment metrics (rank label alignment and silhouette rank alignment). The combination of strong boundary determination and comparatively poorer label alignment suggests that, while CSOA effectively distinguishes a cell subpopulation expressing a given biological signal, it is less well suited for evaluating the strength of that signal within the identified subpopulation. Importantly, CSOA scores represent a relative strength of a biological signal rather than the probability that a cell belongs to a specific biological state. Collectively, the characteristics of CSOA enable it to robustly identify biological signals and outperform existing approaches, thereby advancing single-cell gene set enrichment analysis across diverse applications, including tissue development, regeneration, and disease research.

## Key points

– CSOA is a novel scGSEA method that leverages pairwise gene relationships to compute gene set scores.
– CSOA employs overlaps of pairs of high-expression cell sets built for each signature gene, assessing the statistical significance of these overlaps, then ranking, filtering, and scoring these overlaps to calculate per-cell scores.
– The study compares CSOA with 16 existing methods and introduces a novel benchmarking framework.

## Data Availability

CSOA is available at https://bioconductor.org/packages/release/bioc/html/CSOA.html and https://github.com/andrei-stoica26/CSOA. The code used to generate the results in this paper is available at https://github.com/andrei-stoica26/CSOAResults. The package used for benchmarking, GSABenchmark, is available at https://bioconductor.org/packages/release/bioc/html/GSABenchmark.html and https://github.com/andrei-stoica26/GSABenchmark.

---

**Figure S1:**
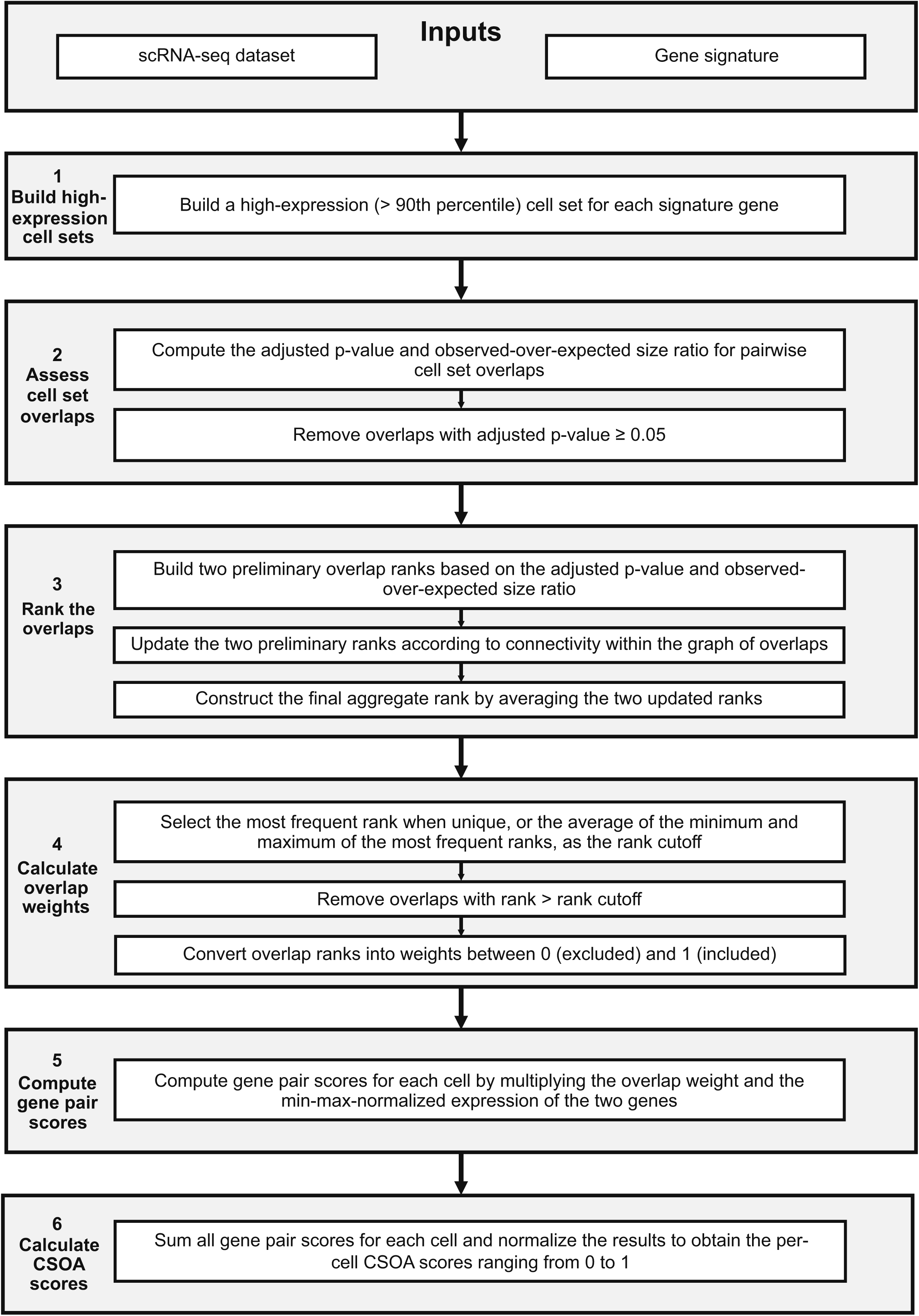
Workflow of CSOA. CSOA uses a single-cell RNA-seq dataset and a gene set as inputs to calculate pairwise overlaps among high-expression cell sets and derive per-cell gene set scores.

**Figure S2:**
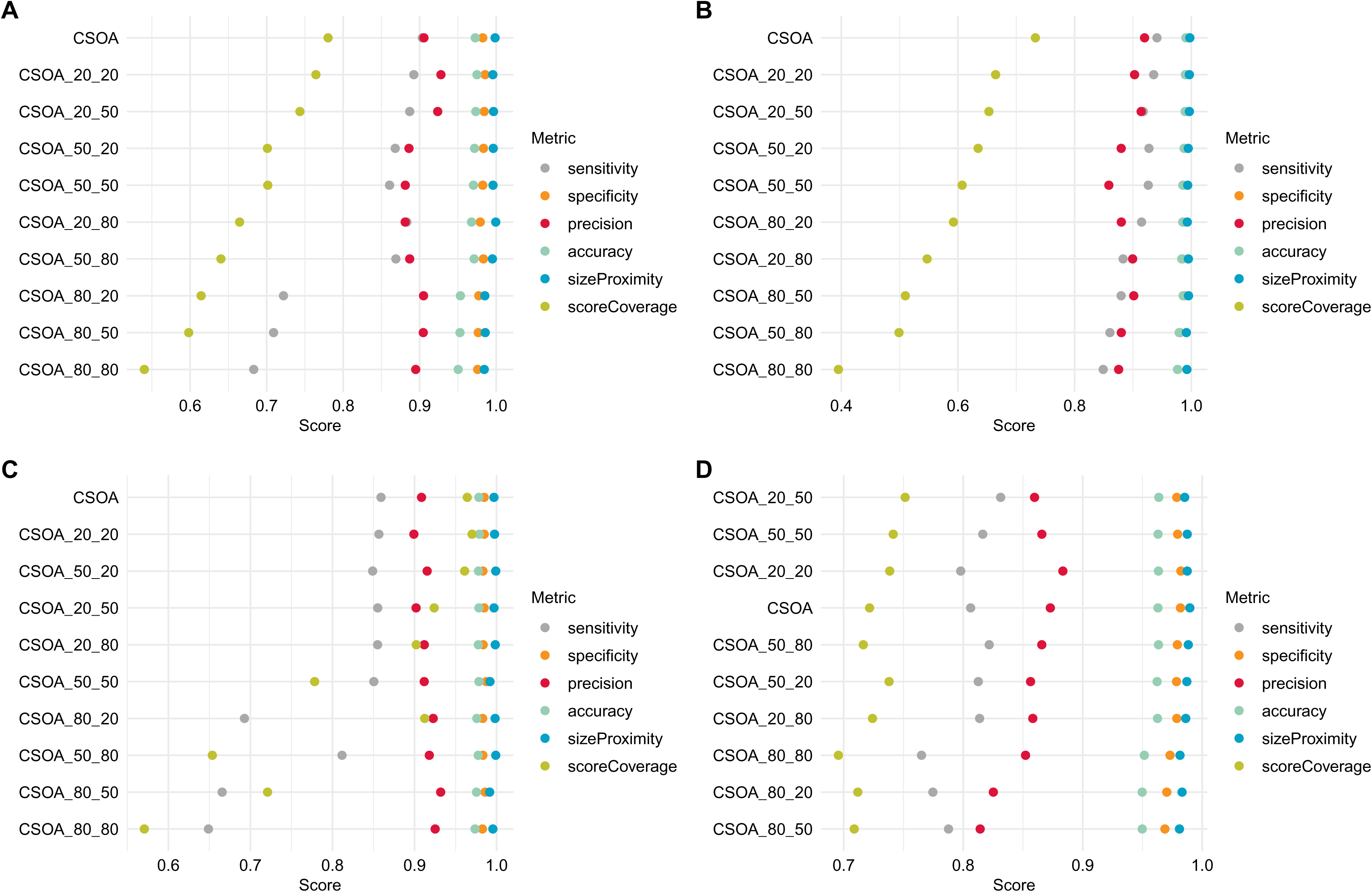
Assessment of class boundary determination robustness to gene loss and noise. In the class boundary determination benchmark, CSOA retained good performance for most choices of gene loss and noise on the Baron pancreas (A), lung proximal airway stromal cells (B), Merkel cell carcinoma (C) and PBMC (D) datasets.

**Figure S3:**
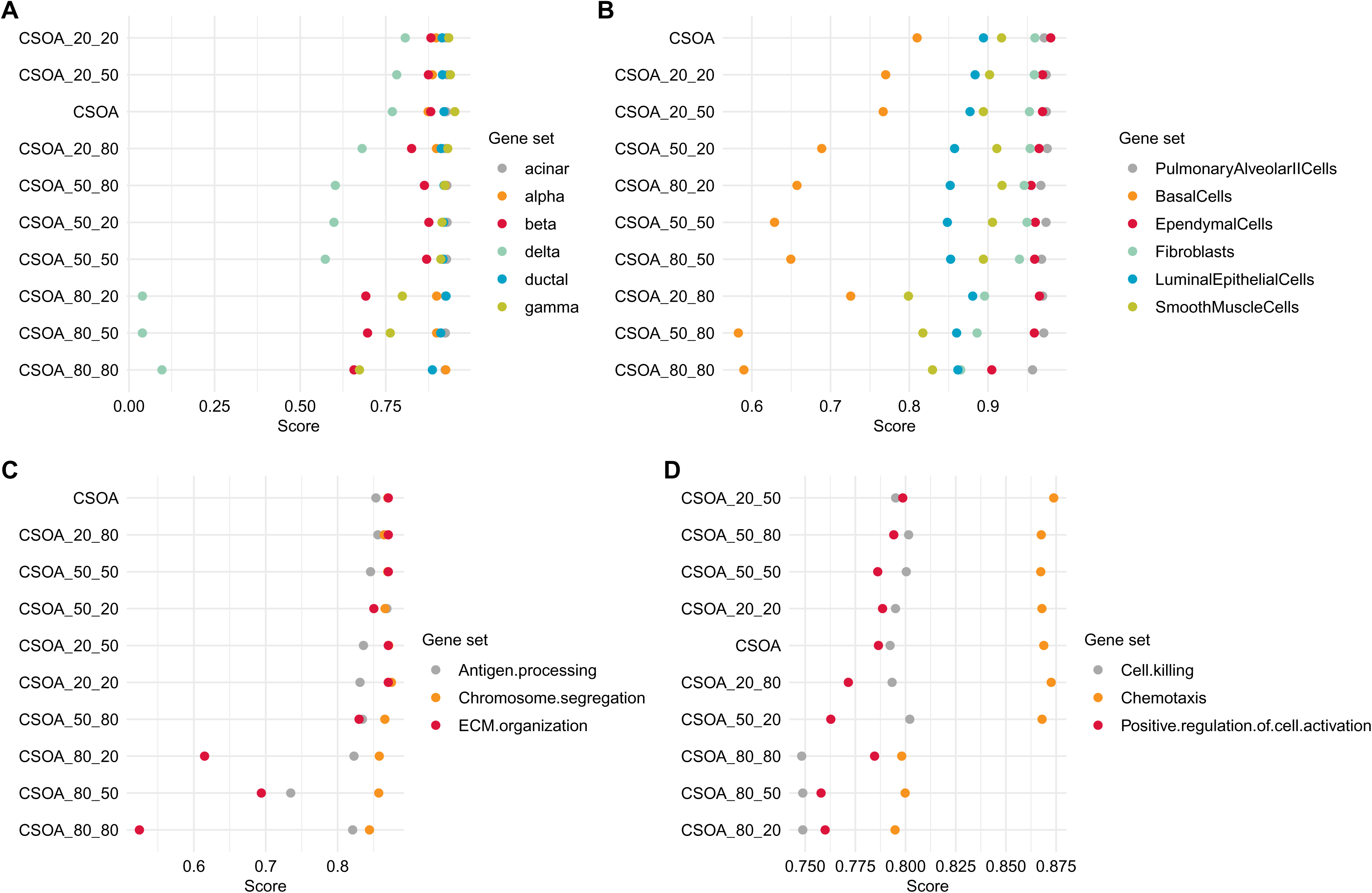
Assessment of comprehensive MCC robustness to gene loss and noise. In the comprehensive MCC assessment, CSOA retained good performance for most choices of gene loss and noise on the Baron pancreas (A), Merkel cell carcinoma (C) and PBMC (D) datasets, while declines in performance associated with a gene loss of 50% or higher are observed on the lung proximal airway stromal cells (B).

**Figure S4:**
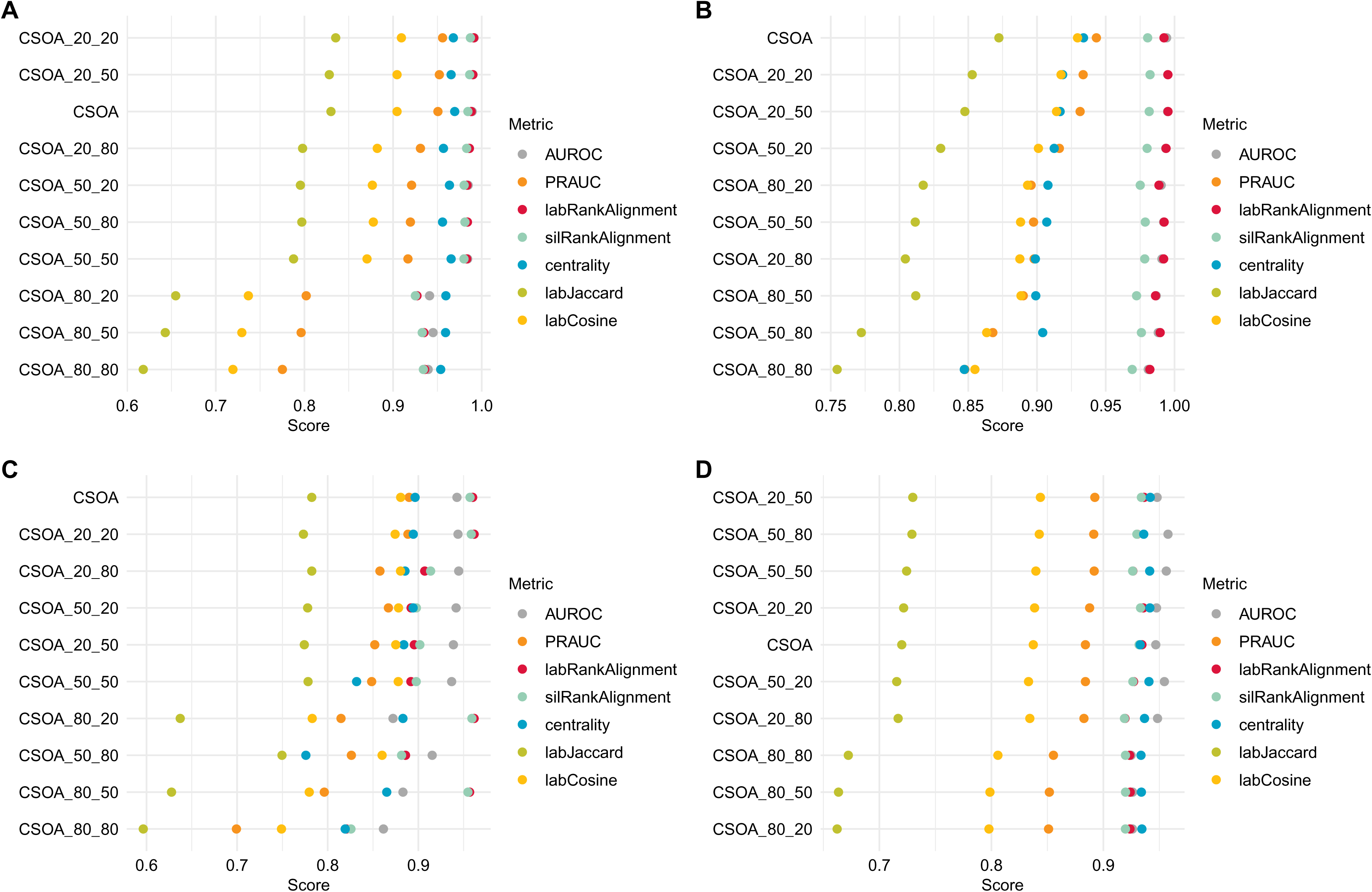
Global evaluation robustness to gene loss and noise. In the global evaluation benchmark, CSOA retained good performance for most choices of gene loss and noise on the Baron pancreas (A), lung proximal airway stromal cells (B), Merkel cell carcinoma (C) and PBMC (D) datasets.

**Figure S5:**
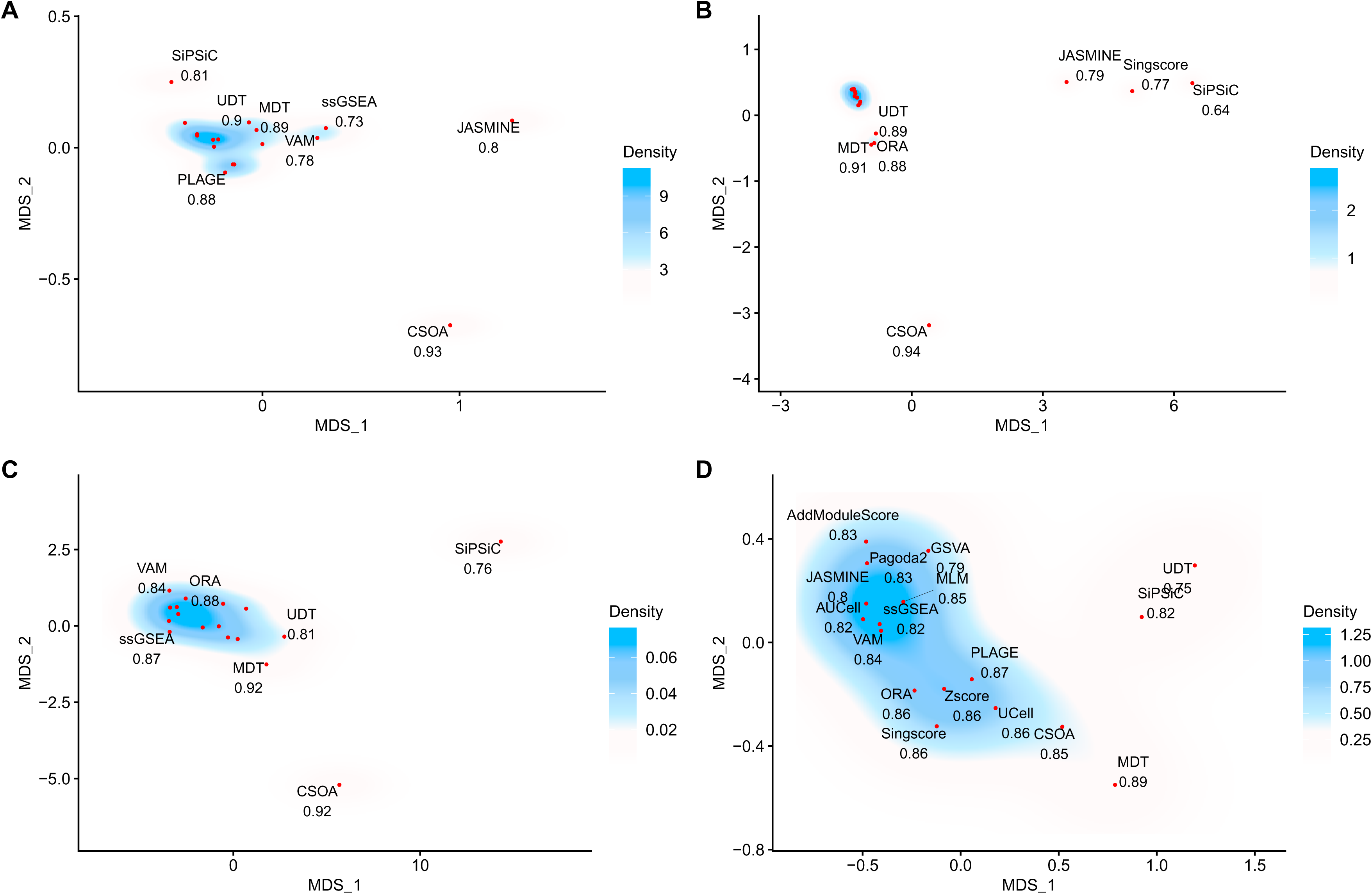
Method similarity analysis based on multidimensional scaling (MDS) of scores-derived percentages. When scores converted to percentages of the total score were assessed, CSOA was highly divergent from the other methods on the Baron pancreas (A), lung proximal airway stromal cells (B) and Merkel cell carcinoma (C) datasets, and relatively divergent on the PBMC dataset (D).

**Figure S6:**
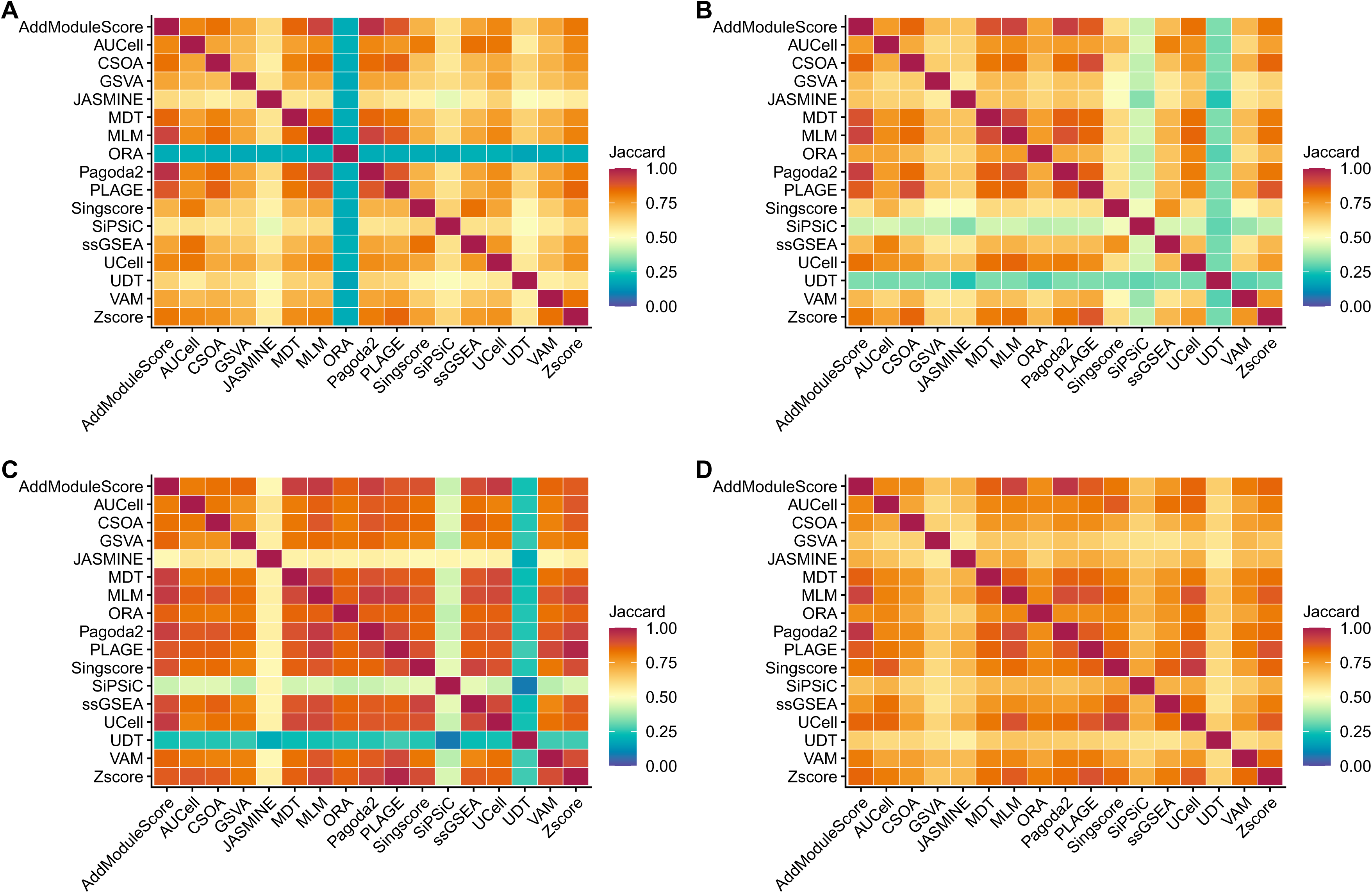
Method similarity analysis based on Jaccard scores of binary predictions. When binary predictions were assessed, CSOA delivered results broadly similar to those of other high-ranking methods such as PLAGE on the Baron pancreas (A), lung proximal airway stromal cells (B), Merkel cell carcinoma (C), and PBMC (D) datasets.

**Figure S7:**
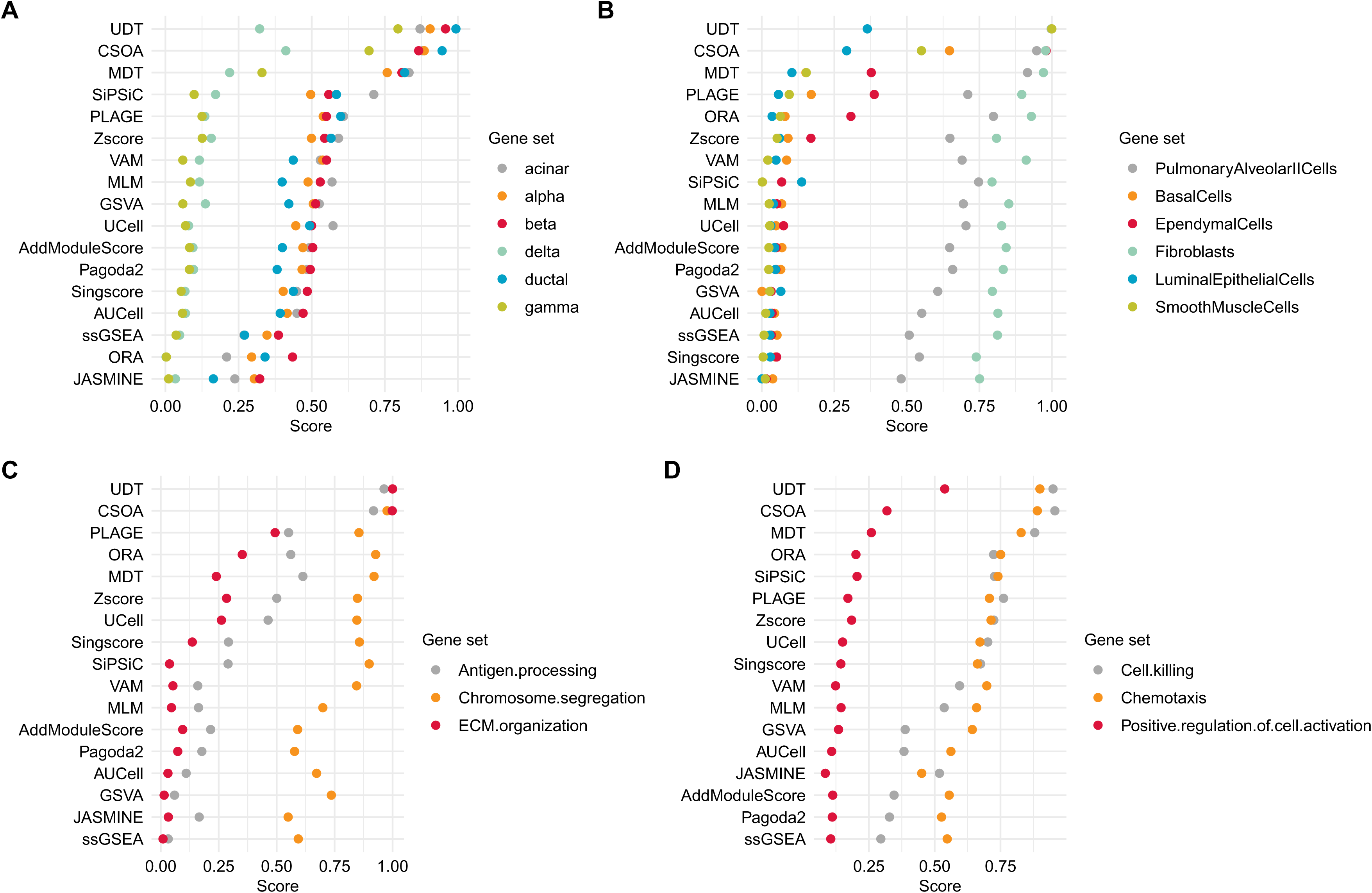
Score coverage benchmark. In terms of score coverage, CSOA ranked second on the Baron pancreas (A), lung proximal airway stromal cells (B), Merkel cell carcinoma (C), and PBMC (D) datasets.

**Figure S8:**
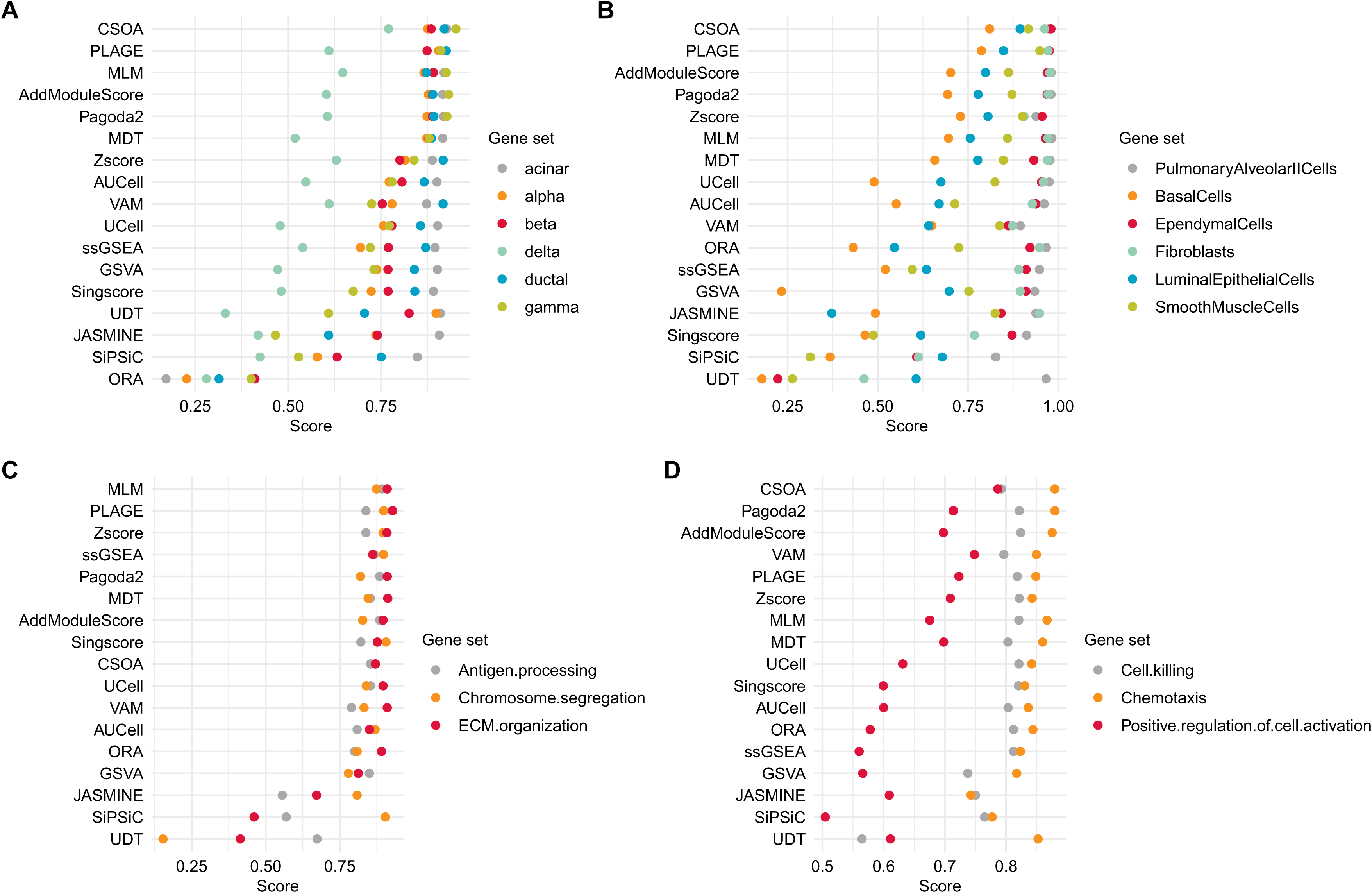
Direct MCC benchmark. In the direct MCC assessment, CSOA ranked the first on the Baron pancreas (A), lung proximal airway stromal cells (B), and PBMC (D) datasets, and the ninth on the Merkel cell carcinoma dataset (C).

**Figure S9:**
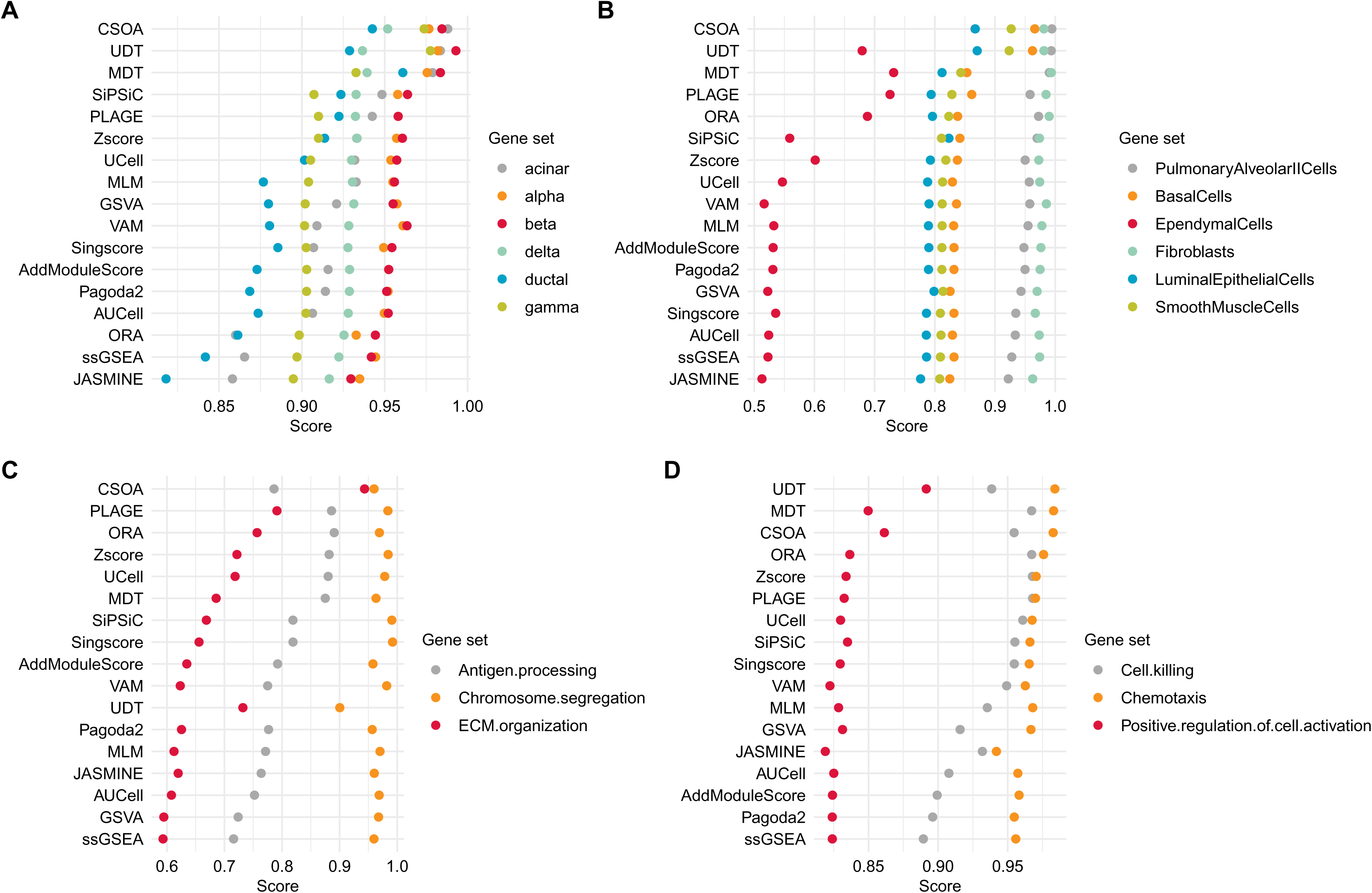
Centrality benchmark. In terms of centrality, CSOA ranked the first on the Baron pancreas (A), lung proximal airway stromal cells (B), and Merkel cell carcinoma datasets (C), and the third on the PBMC dataset (D).

## Tables

**Table S1.** The human scRNA-seq datasets used to benchmark CSOA.

| Dataset | Number of cells | Source | Usage |
| --- | --- | --- | --- |
| Baron pancreas dataset | 8,569 | The scRNAseq R package | Identifying cell types |
| Lung proximal airway stromal cells | 11,186 | PanglaoDB<br>(SRA640325_SRS2769051) | Identifying cell types |
| Merkel cell carcinoma cells | 5,627 | PanglaoDB<br>(SRA749327_SRS3693909) | Identifying functional identities |
| Peripheral blood mononuclear cells | 10,940 | PanglaoDB<br>(SRA550660_SRS2089639) | Identifying functional identities |

